# Intensity of infection with intracellular *Eimeria* spp. and pinworms is reduced in hybrid mice compared to parental subspecies

**DOI:** 10.1101/683698

**Authors:** Alice Balard, Víctor Hugo Jarquín-Díaz, Jenny Jost, Iva Martincová, Ľudovít Ďureje, Jaroslav Piálek, Miloš Macholán, Joëlle Goüy de Bellocq, Stuart J. E. Baird, Emanuel Heitlinger

## Abstract

The longstanding impression that hybrid mice are more highly parasitized and therefore less fit than parentals persists despite the findings of recent studies. Working across a novel transect of the European House Mouse hybrid zone we assessed intracellular infections by *Eimeria*, a parasite of high pathogenicity, and infections by pinworms, assumed to be less pathogenic. For *Eimeria* we found lower intensities in hybrid hosts than in parental mice but no evidence of lowered probability of infection in the centre of the hybrid zone. This means ecological and epidemiological factors are very unlikely to be responsible for the reduced load of infected hybrids. Focussing on parasite intensity (load in infected hosts) we also corroborated reduced pinworm loads reported for hybrid mice in previous studies. In addition we questioned whether differences in body condition during infection would indicate different impacts on hybrid vs. parental hosts’ health. We couldn’t show such an effect. We conclude that intensity of diverse parasites, including the previously unstudied *Eimeria*, is reduced in hybrid mice compared to parental subspecies. We suggest caution in extrapolating this to differences in hybrid host fitness in the absence of, for example, evidence for a link between parasitemia and health.

## Introduction

Hybrid zones can be studied over decades and allow inference regarding the impact of the different endogenous and exogenous forces at play in the process of hybridization (Barton and Hewitt 1985). The European house mouse hybrid zone (HMHZ) is a tension zone characterized by selection against hybrids replaced by immigrating less admixed mice (Barton and Hewitt 1985). After ∼500 000 years of (mostly) allopatric divergence two house mouse subspecies, *Mus musculus domesticus* and *Mus musculus musculus* (hereafter Mmd and Mmm), have come into secondary contact in Europe as a result of different colonization routes south and north of the Black Sea, respectively (Boursot et al. 1993, Duvaux et al. 2011). The HMHZ is about 20 km wide and more than 2500 km long, running from Scandinavia to the coast of the Black Sea (Boursot et al. 1993, Macholán et al. 2003, Jones et al. 2010, Baird and Macholán 2012). This zone represents a semi-permeable barrier to gene flow between the two taxa (Macholán et al. 2007, 2011). The main selective forces acting against hybrids are thought to be endogenous rather than ecological (Boursot et al. 1993, Baird and Macholán 2012), for example disruption of spermatogenesis in hybrids (Albrechtová et al. 2012, Turner et al. 2012).

The relevance of hybridization, producing individuals admixed between genetically distinct populations, is increasingly recognized by biologists. Mallet (2005) suggested that hybridization occurs in more than 10% of animal species and 25% of vascular plant species. Recently, the realization that our own species is a product of hybridization has raised interest further (Green et al. 2010). In a conservation context hybridization with introduced species can threaten autochthonous endangered animals (Simberloff 1996). Parasites are omnipresent in natural systems and so it is important for biologists interested in hybridization to comprehend the interplay between parasites and hosts under hybridization.

The HMHZ was one of the first animal hybrid zones studied for differences in parasite loads (Sage et al. 1986). Parasites are traditionally seen as decreasing their hosts’ fitness, and differences in resistance to parasites between hybrid and pure hosts was suggested to affect the dynamics of hybrid zones (Fritz et al. 1999). This traditional framework postulates differences in parasite loads in hybrids vs. parental hosts to result in effects on the strength of host species barriers.

Initial results on parasites obtained in the HMHZ and experimental studies seemed to indicate elevated parasite loads in hybrids. This has been interpreted as potentially leading to fitness reductions in hybrids, hampering hybridization and thus reinforcing species barriers (Sage et al. 1986, Moulia et al. 1991, Moulia et al. 1993). Infection experiments using the protozoan *Sarcocystis muris* led to a similar conclusion (Derothe et al. 2001). Other laboratory experiments, however, showed either no hybrid effect on helminth load or even reduced load in hybrids compared to pure mouse strains (Moulia et al. 1995, Derothe et al. 2004). The field study with arguably the highest statistical power found reduced helminth loads (especially the pinworms *Aspiculuris tetraptera* and *Syphacia obvelata* and the whipworm *Trichuris muris*) in hybrid mice (Baird et al. 2012). It should also be noted that the design of the field studies preceding the Baird et al. (2012) reappraisal usually suffered from low sample sizes and/or maintenance of mice under laboratory conditions before assessment of parasite burden, which may have allowed spurious signal to dominate the results. Nevertheless, even the basic direction of parasite load differences in hybrid mice compared to parental genotypes seems still controversial to some researchers.

We now see that, despite working within the framework of the same hybrid zone, two different interpretations of parasite loads in hybrid mice have arisen. One way forward in such circumstances is to check over replicates. To distinguish between the load interpretations we therefore, in a new transect replicate of the HMHZ, asked if (1) parasite loads are higher or lower in hybrids compared to parentals, and (2) if these loads are consistent, or differ, across two levels of pathogenicity.

Pinworms (oxyurids) have been shown to be the most prevalent helminths infecting house mice in the HMHZ (Goüy de Bellocq et al. 2012). They are broadly distributed, can reinfect their hosts throughout their lives, and may be considered almost non-pathogenic (Taffs 1976). *Eimeria* spp. are often considered host-specific, with several thousand species parasitizing different vertebrates (Haberkorn 1970, Chapman et al. 2013). These parasites infect the intestinal epithelial cells of vertebrates and induce symptoms such as weight loss and diarrhoea. For example, infecting the NMRI mouse laboratory strain with *Eimeria* oocysts isolated from mice captured in the HMHZ resulted in a weight loss up to 20% compared to control (Al-khlifeh et al. 2019). Work in wild rodents indicates high pathogenicity of *Eimeria* spp. under field conditions: in populations of bank voles (*Myodes glareolus*), *Eimeria* spp. reduce body condition of both mothers and of offspring at birth (Hakkarainen et al. 2007). In deer mice these *Eimeria* affect overwinter survival of males (Fuller and Blaustein 1996). In the European HMHZ, three *Eimeria* species have been identified: *E. ferrisi*, *E. falciformis*, and *E. vermiformis* with prevalences of 16.1%, 4.2% and 1.1%, respectively (Jarquín-Díaz et al. 2019).

We assessed *Eimeria* infection in a novel transect of the HMHZ in Brandenburg, northeastern Germany, testing the impact of host hybridization on intensity of this parasite. By focussing on parasite intensity (extent of parasite infection in only infected hosts; Bush et al. 1997), we arguably exclude ecological and epidemiological factors for differences in load (i.e. parasite prevalence and abundance, the latter defined as parasite load in all hosts). We show that (1) parasite loads are consistently lower in hybrids compared to parental genotypes in the HMHZ and (2) that this pattern is consistent across two different levels of pathogenicity.

## Material & Methods

### Sampling

Our sampled individuals consist of 660 house mice trapped using live traps placed in farms or houses between 2014 and 2017. The study area ranges from 51.68 to 53.29 degrees of latitude (200 km) and from 12.52 to 14.32 degrees of longitude (140 km). Each year mice were trapped in September when it is possible to capture a high number of mice in this region. In addition, sampling at the same season every year reduces potential seasonal variation (Haukisalmi et al. 1988, Abu-Madi et al. 2000). The trapping was designed to capture both parental and hybrid/recombinant populations. Mice were individually isolated in cages and then euthanized by isoflurane inhalation followed by cervical dislocation and dissection within 24 hours after capture (animal experiment permit No. 2347/35/2014). Tissue samples (muscle and spleen) were put to liquid nitrogen and stored at −80°C for subsequent host genotyping. Digestive tracts were dissected and inspected for helminth parasites (see below). Ileum, caecum and colon tissues were frozen in liquid nitrogen and then stored separately at −80°C. Individual mice were measured (body length from nose to annus) and weighted.

### Host genotyping

The admixture proportion of mouse genomes across the HMHZ was estimated for each mouse as a value of the hybrid index (HI) calculated as a proportion of Mmm alleles in a set of 4-14 diagnostic markers (at least 10 loci in 92% of the mice). This set consists of one mitochondrial marker (*Bam*HI, a restriction site in the *Nd1* gene; Munclinger et al. 2002, Božíková et al. 2005), one Y-linked marker (presence/absence of a short insertion in the *Zfy2* gene; Nagamine et al. 1992, Boissinot and Boursot 1997), six X-linked markers (three B1 and B2 short interspersed nuclear elements in *Btk*, *Tsx* (Munclinger et al. 2003), and *Syap1* (Macholán et al. 2007), *X332*, *X347* and *X65* (Dufková et al. 2011, Ďureje et al. 2012)), and six autosomal markers (*Es1*, *H6pd*, *Idh1*, *Mpi*, *Np*, *Sod1*; Macholán et al. 2007). HIs ranged from 0 to 1, HI of 0 indicating a pure Mmd and HI of 1 a pure Mmm (Macholán et al. 2007, Baird et al. 2012).

The course of the HMHZ across the study area was estimated using the program Geneland v4.0.8 (graphical resolution increased over defaults) based on a subset of the six autosomal markers that were genotyped in all mice. Geneland uses a Markov chain Monte Carlo (MCMC) approach to combine both geographical and genetic information (Guillot et al. 2005). The number of clusters was set to 2, 10^6^ MCMC iterations were performed and saved every 100th iterations (10^4^ iterations saved). The first 200 iterations were discarded as burn-in and the resolution of the map was set to 2000 pixels for the x axis and 1400 for the y axes corresponding roughly to 1 pixel for 100m (Macholán et al. 2011).

### Parasite load estimation

Mouse digestive tracts were dissected and inspected for helminth presence with a binocular microscope. Helminths were counted and stored in 70% ethanol for later identification by molecular analysis and, when more than one worm per host was present, in 3.5% formalin for later morphological comparison with species descriptions. In this study we considered only the most prevalent helminths, the oxyurids *Syphacia obvelata* and *Aspiculuris tetraptera*.

DNA was extracted from ileum and caecum tissues and quantitative PCR (qPCR) was used for estimation of *Eimeria* spp. load. DNA extraction was performed using the innuPREP DNA Mini Kit (Analytik Jena AG, Jena, Germany) following the instructions of the manufacturer with additional mechanical tissue disruption with liquid nitrogen in a mortar. Both quality and quantity of isolated DNA were measured by spectrophotometry in a NanoDrop 2000c (Thermo Scientific, Waltham, USA). The presence of *Eimeria* spp. was tested using qPCR to detect intracellular stages of the parasite as well as house mouse house keeping gene as internal reference. Primers used for *Eimeria* spp. detection targeted a short mitochondrial *COI* region (Eim_COI_qX-F: TGTCTATTCACTTGGGCTATTGT; Eim_COI_qX-R: GGATCACCGTTAAATGAGGCA), while *Mus musculus* primers targeted the *CDC42* nuclear gene (Ms_gDNA_CDC42_F: CTCTCCTCCCCTCTGTCTTG; Ms_gDNA_CDC42_R: TCCTTTTGGGTTGAGTTTCC). Reactions were performed using 1X iTaqTM Universal SYBR® Green Supermix (Bio-Rad Laboratories GmbH, München, Germany), 400 nM of each primer and 50 ng of DNA template in 20 µL final volume. Cycling amplification was carried out in a Mastercycler® RealPlex 2 thermocycler (Eppendorf, Hamburg, Germany) with the following amplification program: 95°C initial denaturation (2 min) followed by 40 cycles of 95°C denaturation (15 s), 55°C annealing (15 s) and 68°C extension (20 s). Melting curve analyses were performed in order to detect primer dimer formation and unspecific amplification. ΔCt was calculated as difference of the threshold cycle (Ct) between mouse and *Eimeria* spp. values (corresponding to a log2 ratio between parasite and mouse DNA). We considered ΔCt = -5 our limit of detection (Ahmed et al. 2019, Jarquín-Díaz et al. 2019). Samples with a ΔCt lower than -5 were considered negative (unspecific signal due to amplification of non-target DNA). Samples with a ΔCt higher than -5 for at least one of the two intestinal tissues were considered positive, and in the case of detection in both tissues, the higher value was taken as a proxy of individual parasite load. This parasite load of the intestinal tissue stage is denoted as “ΔCt_Mouse–Eimeria_” throughout this paper.

### General parasite assessment

As the distributions of parasite loads are expected to be highly skewed (Bliss and Fisher 1953), the median (as an estimator for the mode) is more informative than the mean (Rózsa et al. 2000). We therefore took the median of parasite load across all hosts (median abundance) and of parasite load of infected host (median intensity) for pinworms, and only median intensity for *Eimeria* spp. Prevalence (relative frequency of infected individuals amongst all tested individuals) confidence intervals were obtained with Sterne’s exact method (Sterne 1954, Reiczigel et al. 2010). Calculations were performed using the epiR package (Nunes et al. 2018) running within the R statistical computing environment (R Development Core Team 2008).

### Statistical prediction of probability of infection by parasites along the hybrid zone

The parasite infection process can be split into two components: probability of infection, and parasite burden following infection. Absence of parasite in a given host can result from either absence of exposure to parasite or complete host resistance, while quantitative parasite load depends on intrinsic host or parasite components or their interactions. These two components rely on different mechanisms, and therefore should be assessed separately (Poulin 2013).

Firstly, we considered the predicted probability of infection along the HI. We used a proxy of “genetic distance to zone centre”: for individuals with HI between 0 and 0.5 the proxy is HI, for individuals with HI between 0.5 and 1 the proxy is 1 – HI. This was used to model a dichotomous response variable (uninfected = 0; infected = 1) by logistic regression, as a linear combination of the predictor variables “genetic distance to zone centre” and “Sex” (including interactions). Analyses were done in R with the function glm from the stats package (R Development Core Team 2008).

### Statistical test of the host hybridization effect on parasite intensity

Secondly, it has been shown that macroparasites tend to aggregate within their hosts, the majority of host carrying no or a low burden, and a minority a high one (Shaw and Dobson 1995). We modelled this distribution of parasite burden in infected hosts as negative binomial. Following the approach of Baird et al. (2012), we tested if hybrid mice had higher or lower parasite burdens than that expected in case of additivity (if the relationship between host parasite load and hybrid index was linear). The hybridization level on each individual was modelled as the degree to which new gene combinations are brought together compared to the pure subspecies. This was estimated from the hybrid index using the function for expected heterozygosity (Baird et al. 2012):

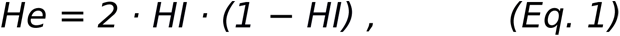

The parasite load for a given HI was then estimated as follows:

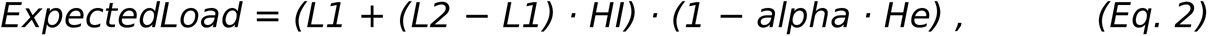

where L1 is the parasite load of pure Mmd, L2 the parasite load of pure Mmm, and alpha the hybridization effect (deviation of parasite estimated load from the additive model). We considered four nested hypotheses increasing in complexity, and compared them with the G-test (likelihood ratio test) to consider a more complex hypothesis only when justified by a significant increase in likelihood. Expected parasite load is fixed to be identical for both subspecies and both host sexes in hypothesis H0. The more complex H1 allows load differences for the host sexes, H2 allows different loads between the subspecies at the extremes of the hybrid index, and H3 allows differences both between the subspecies and sexes.

Adequate distributions of values for each parasite and detection method considered were selected using log likelihood and AIC criteria and by comparing goodness-of-fits plots (density, CDF, Q-Q, P-P) (R packages MASS (Venables and Ripley 2002) and fitdistrplus (Delignette-Muller and Dutang 2015)). The negative binomial distribution should perform well for macroparasite counts (Crofton 1971, Shaw and Dobson 1995), which was confirmed for helminths (Baird et al. 2012). Values of (ΔCt_Mouse–Eimeria_) were found to be well described by the Weibull distribution after being positively shifted.

The Negative Binomial distribution is parameterized by two arguments: its expectation (Expected Load, Eq. 2), and the inverse of its aggregation defined, which is allowed to vary across HI as:

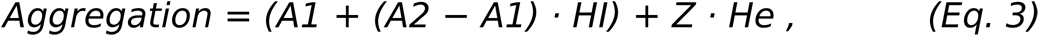

Z being the deviation from the additive model, in proportion to *He*, which is maximal in the zone centre (Baird et al. 2012). The Weibull distribution is parametrized by its shape and scale parameters (allowed to vary freely during maximum likelihood search) linked by the formula:

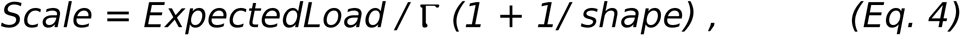

┌ being the gamma function.

We fit the models using likelihood maximisation (using the R package mle2; Bolker and Team 2017). Parasite load was estimated either including or excluding the hybridization effect parameter (by setting HI = 0 in *ExpectedLoad*), and we compared these two models using the G-test. In the case of ΔCt_Mouse–Eimeria_, the Weibull distribution requires positive values as input. Therefore, we estimated an extra “shift parameter” by maximum likelihood at 7.14.

### Test of body condition differences between infected and non-infected mice along the hybrid zone

Residuals from ordinary least squares regression of body weight by body length were estimated for each individual, separately for males and females. Pregnant females were excluded from the analysis. Individuals with a positive residual were considered in better condition than individuals with a negative one, as this index correlates with variation in fat, water, and lean dry mass (Schulte-Hostedde et al 2005). We tested if hybrid mice had higher or lower residuals than that expected for intermediate between pure hybridizing taxa (“additivity”), and if the potential hybridization effect was different between infected and not infected mice, for *Eimeria* spp. as well as for pinworm infections. Differences between the subspecies were allowed.

Values of residuals of body weight by body length regression are well described by the Normal distribution, parametrized by its standard deviation (allowed to vary freely during maximum likelihood searches) and its mean defined as:

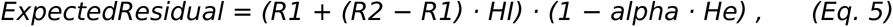

where R1 is the expected residual value of pure Mmd, R2 the expected residual value of pure Mmm, and alpha the hybridization effect.

We fit the models using maximum likelihood (using the R package mle2; Bolker and Team 2017), either including or excluding the hybridization effect parameter (by setting HI = 0 in *ExpectedResiduals*), and we compared these two models using the G-test.

All graphics were produced using the R packages ggplot2 (Wickham 2016) and ggmap (Kahle and Wickham 2013), and compiled using the free software inkscape v.0.92 (https://inkscape.org).

## Results

### Host genotyping and characterization of the HMHZ for a novel transect

We caught and genotyped a total of 660 mice (363 females, 297 males) over four sampling seasons (2014: *N*=87; 2015: *N*=163; 2016: *N*=167; 2017: *N*=243) at 154 localities. A median of 2 mice per locality were captured. A list of individual hybrid indices, georeferences, and parasite loads is available in supplementary table S1. As shown in Fig. 1, the HMHZ runs across the former East Germany, making a broad arc around the city of Berlin, approaching within ca. 20 km of the bordering Oder River near Eberswalde.

**Figure 1.**
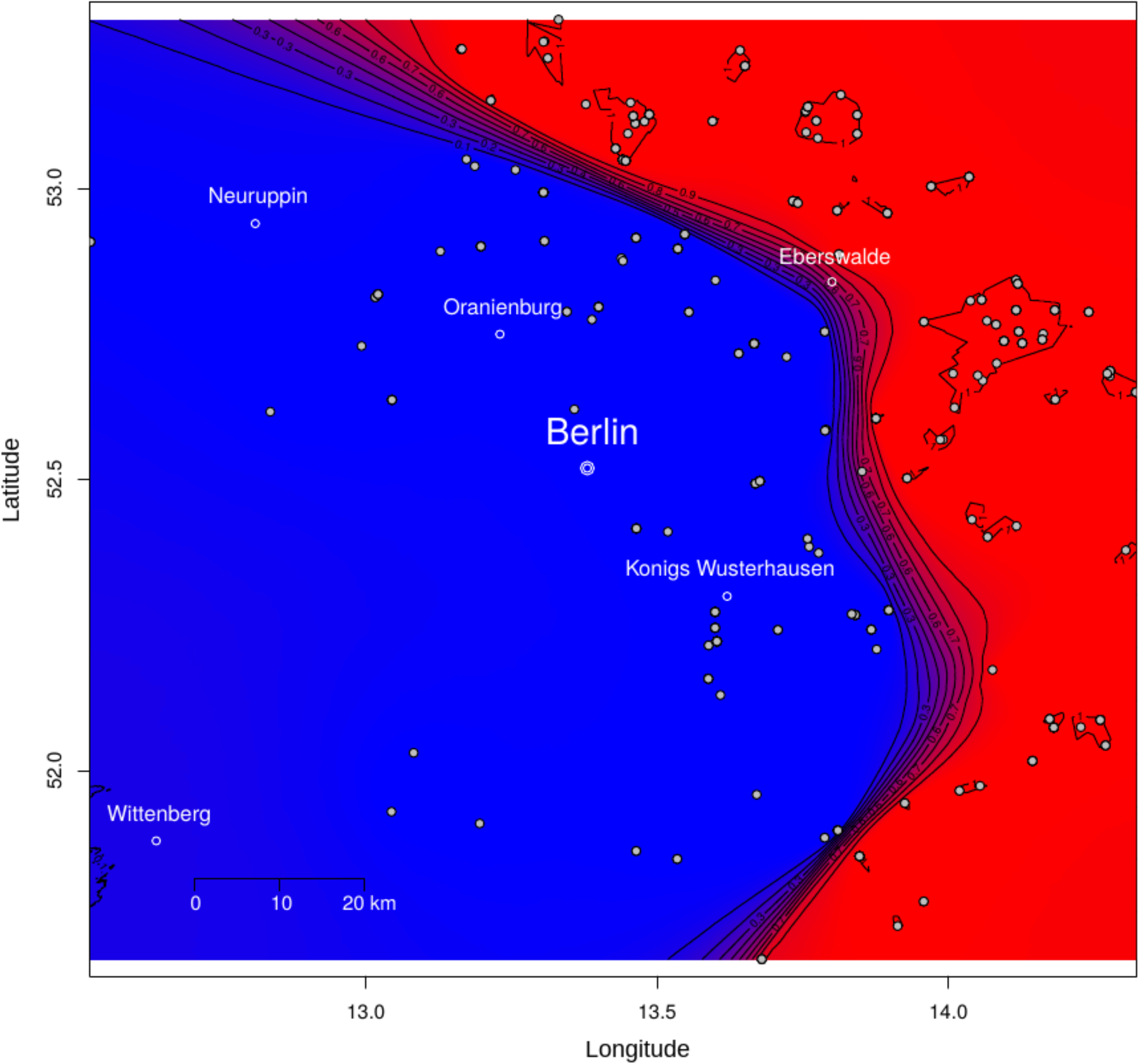
Geographic range of house mouse subspecies in the European house mouse hybrid zone. Spatial organization of the HMHZ was inferred using six autosomal markers (*Es1*, *H6pd*, *Idh1*, *Mpi*, *Np*, *Sod1*). *Mus musculus domesticus* is found west of the hybrid zone (blue), *Mus musculus musculus* east of it (red). The numbers at the level contours indicate posterior probabilities of population membership for each mouse subspecies. White dots represent each mouse included in the study.

### Parasite prevalence and intensity

To investigate *Eimeria* infections we checked 384 mice sampled in 2016 and 2017 for the presence and intensity of tissue stages (Fig. 2a). The estimated parasite prevalence was 18.2% (70/384) (Sterne’s Exact method CI 95%: [14.5, 22.5]). To quantify the intensity of infection we determined the amount of *Eimeria* mitochondrial DNA per host nuclear DNA using ΔCt_Mouse–Eimeria_. The median *Eimeria* intensity was -2.4 corresponding to 5.2 times less parasite mitochondrial DNA than host nuclear DNA.

**Figure 2.**
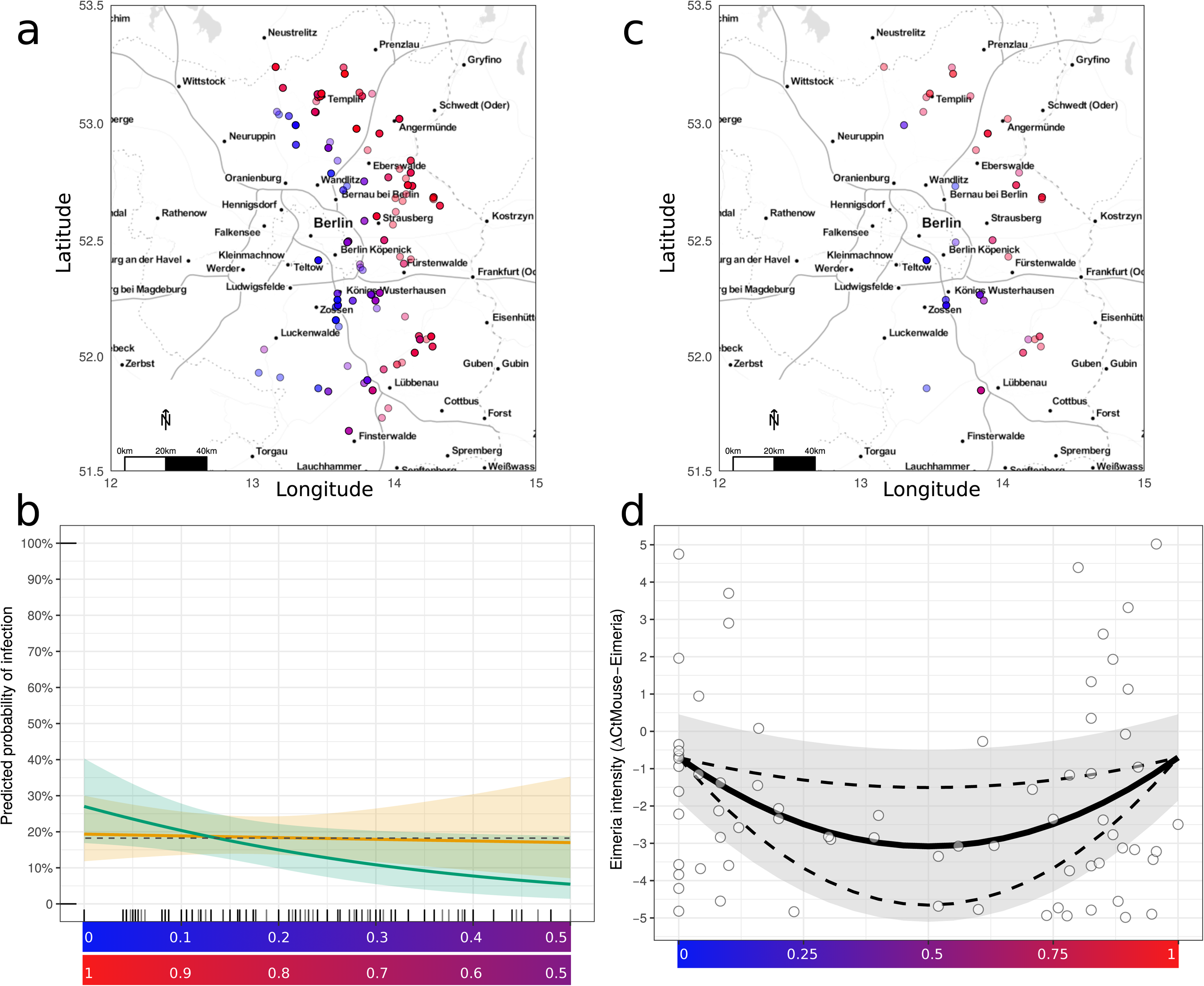
Probability of infection is constant and intensity of *Eimeria* infection is reduced in hybrids. Individual mice tested for detection and quantification of *Eimeria* spp. tissue stages (a) and mice tested positive (c) are displayed on a map. The predicted probability of infection does not differ in more admixed mice (b) for males (green) and females (orange)(average observed probability of infection: grey dotted line). *Eimeria* intensity is reduced at intermediate values of the hybrid index (d), represented as a gradient ranging from 0 (pure Mmd, in blue) to 1 (pure Mmm, in red). The optimized fit is represented by a solid line, the 95%CI of the fit as all parameters are allowed to vary in their 95%CI, is plotted as a grey ribbon. The 95%CI of the hybridization parameter alpha, as all parameters are fixed to their fitted value while alpha is allowed to vary in its 95%CI, is plotted as dashed lines.

Between 2014 and 2017, 585 mice were investigated for helminths (Fig. 3a). Prevalence of pinworms in the transect was 52.5% (307/585) (Sterne’s Exact method CI 95%: [48.4, 56.5]) with a median abundance of 1 pinworm per mouse and median intensity of 13 pinworms per infected mouse (maximum number of pinworms in one host: 489).

**Figure 3.**
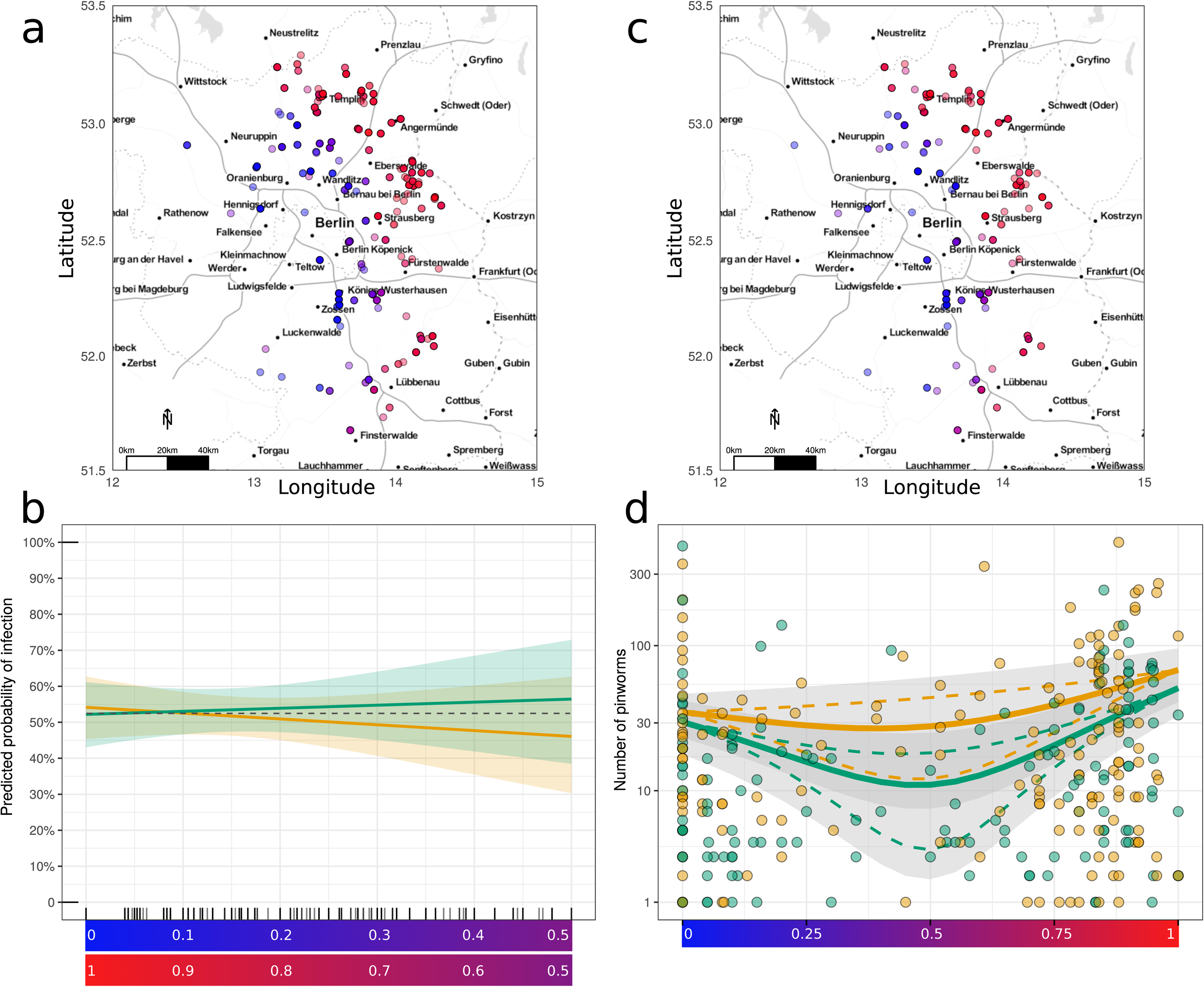
Probability of infection is constant and intensity of pinworm infection is reduced in hybrids. Individual mice tested for detection and quantification of pinworms (a) and mice tested positive (c) are displayed on a map. The predicted probability of infection does not differ in more admixed mice (b) for males (green) and females (orange)(average observed probability of infection: grey dotted line). Pinworm intensity is reduced at intermediate values of the hybrid index (d), represented as a gradient ranging from 0 (pure Mmd, in blue) to 1 (pure Mmm, in red), for males (green) and females (orange). The optimized fit is represented by a solid line, the 95%CI of the fit as all parameters are allowed to vary in their 95%CI, is plotted as a grey ribbon. The 95%CI of the hybridization parameter alpha, while all parameters are fixed to their fitted value and alpha is allowed to vary in its 95%CI, is plotted as dashed lines.

### Similar prevalence of parasites across the zone

In order to control for the simple case of a host density trough at the zone centre, we tested if the probability of being infected was significantly lower for individuals at the host zone centre. Logistic regression using a linear combination of the predictor variables “genetic distance to zone centre” and “Sex” (including interactions) didn’t show any statistically significant effect (p > 0.05) on the probability of infection, neither for *Eimeria* spp. (Fig. 2b) nor for pinworms (Fig. 3b). We therefore could not find evidence of significantly more or less uninfected hosts in the centre hybrid zone, neither for *Eimeria* nor pinworms.

### *Eimeria* spp. load is lower in infected hybrid vs pure Mmm and Mmd mice

To test more specifically the intrinsic host-parasite interplay of hybrids compared to pure mice, we considered only individuals infected by *Eimeria* spp. tissue stages (*N* = 70). Complex models involving differences between sexes and parental taxa did not fit the data significantly better than the null model (Supplementary Table S2). The fit involving the hybridization effect, however, showed significantly higher likelihood than the model without it (G-test; p-value = 0.02). Infected hybrids had significantly lower load of *Eimeria* spp. tissue stages than expected if the load was linear along the hybrid index, with a hybridization effect parameter alpha of 0.74 (Fig. 2d, values of parameters of the fitted model given in Table 1).

**Table 1.** Parametrisation of fitted models. Parameters estimated by maximum likelihood for each dataset. Alpha is the hybridization effect (deviation of parasite estimated load from the additive model) given with its significance p-value. If sexes are separated, corresponding parameters for each sex are given with symbols ♀ and ♂. Nested hypotheses are as follow. H0: same expected load for the subspecies and between sexes; H1: same expected load across sexes, but can differ across subspecies; H2: same expected load across subspecies, but can differ between the sexes; H3: expected load can differ both across subspecies and between sexes. *Mus musculus domesticus* and *Mus musculus musculus* are named hereafter Mmd and Mmm.

| ***Eimeria***<br>**intensity** | Hyp. | Alpha<br>(p-value) | Load in $\Delta C_t$ for both<br>parental subspecies | Shape |
| --- | --- | --- | --- | --- |
| present study | H0 | 0.74 (0.02) | -0.70 | 2.33 |

| **Pinworm**<br>**intensity** | Hyp. | Alpha<br>(p-value) | Load in count<br>Mmd | Load in count<br>Mmm | Aggregation<br>Mmd | Aggregation<br>Mmm | Z parameter |
| --- | --- | --- | --- | --- | --- | --- | --- |
| present study | H3 | ♀ 0.91<br>(0.04) | ♀ 35.57 | ♀ 68.67 | ♀ 1.45 | ♀ 2.00 | ♀ -1.04 |
|  |  | ♂ 1.46<br>(<0.001) | ♂ 30.38 | ♂ 51.86 | ♂ 2.10 | ♂ 1.33 | ♂ -1.23 |
| present study<br>(data from Baird<br>et al. 2012) | H1 | 1.21 (<0.001) | 94.37 | 46.81 | 1.88 | 1.34 | -0.13 |

### Pinworm load is lower in infected hybrid vs. pure Mmm and Mmd mice

We tested pinworm intensity (*N* = 307) in infected hybrids comparing them to infected ‘pure parental’ mice in our Brandenburg transect, excluding potential ecological and epidemiological confounders in the same way. The model allowing differences between the parental taxa and sexes (H3) was found to fit our observations significantly better than the less complex models (Supplementary Table S2). For both sexes, the fit including the hybridization effect showed significantly higher likelihood than the model without it (G-test; p-value = 0.04 for females, p-value < 0.001 for males). Infected hybrids had significantly lower pinworm load than expected if the load was linear along the hybrid index, with the hybridization effect parameter alpha 0.91 (females) and 1.46 (males) (Figure 3d, values of parameters of the fitted model given in Table 1).

### Comparison of pinworms loads with previous reports

To compare the strength of the hybridization effect between our Brandenburg transect and the Czech-Bavarian portion of the HMHZ we applied the H1 model (differences between the taxa but not between the host sexes) to our pinworm abundance data, once with freely varying alpha (fit 1), and once with alpha set to 1.39 as in Baird et al. (2012) (fit 2). Within fit 1, alpha was found significant (G-test; p-value < 0.001). The comparison between the model with freely varying alpha (fit 1) and that using fixed alpha (fit 2) showed no significant likelihood difference (G-test; p-value = 0.11). Therefore, we can conclude that pinworm load differences found in hybrids in this study are consistent with the results obtained in the previously studied Czech-Bavarian transect (Baird et al. 2012).

### No evidence of body condition differences between infected and non-infected mice along the hybrid zone

To test whether infections have a different effect in hybrids vs. parental mice we assessed body condition, which could be a better proxy for host health than parasite load. Modelling of the residuals from ordinary least squares regression of body weight by body length along the hybrid zone (Fig. 4a) did not show a statistically significant hybridization effect (G-test; p-value > 0.05 in both parasite datasets considered). When infected and non-infected individuals were considered separately, neither *Eimeria* spp. infected individuals (G-test; p-value = 0.58) nor *Eimeria* spp. non-infected individuals (G-test; p-value = 0.90) showed a hybridization effect in body condition index (Fig. 4b). The same was true for pinworm infected individuals (G-test; p-value = 0.44) or pinworm non-infected individuals (G-test; p-value = 0.96; Fig. 4c).

**Figure 4.**
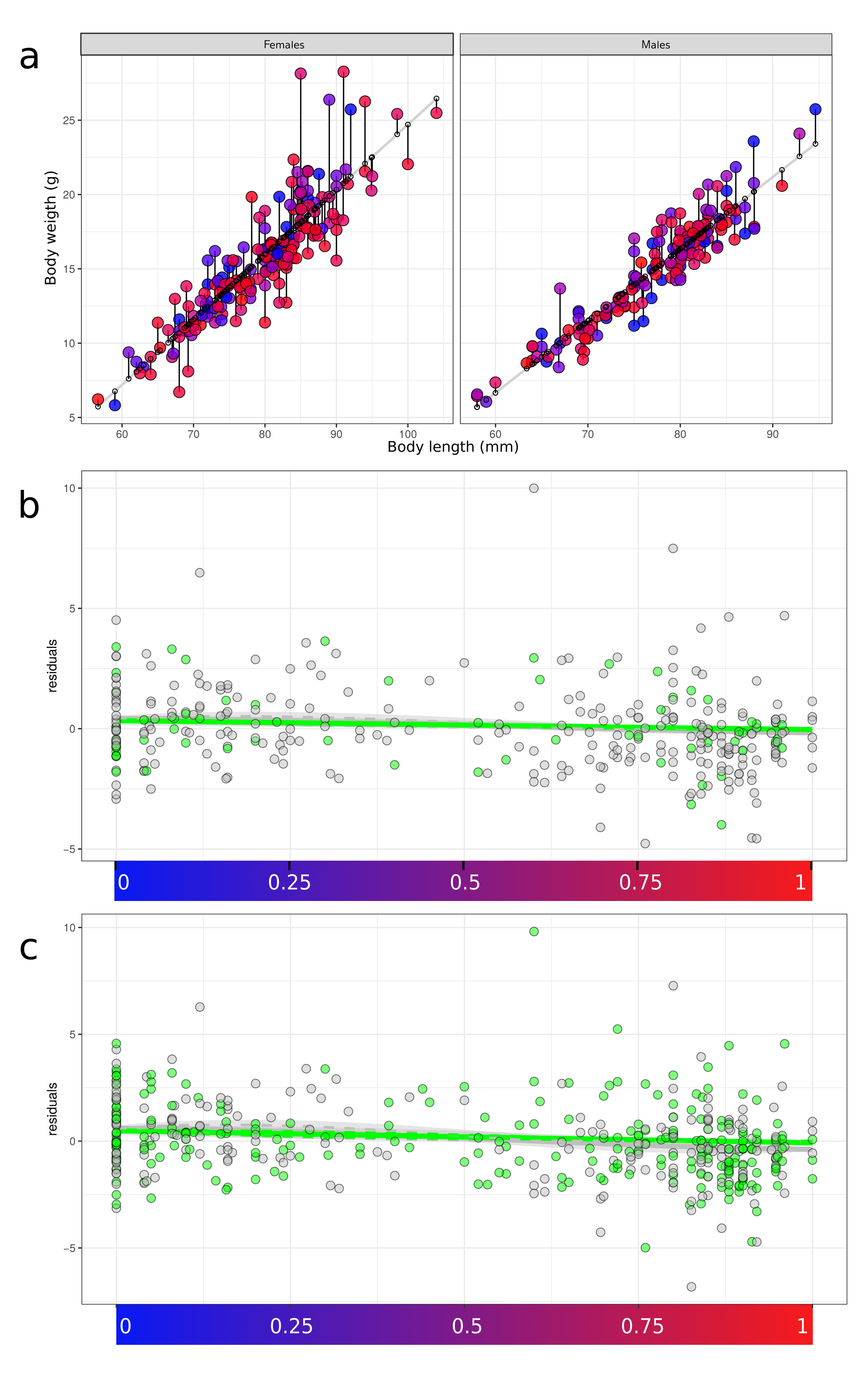
Body condition does not significantly differ between hybrids and pure mice upon infection. We modelled the residuals from ordinary least squares regression of body weight by body length along the hybrid zone. The fit and residuals for female and male mice is given in (a). The hybrid index is represented as a gradient ranging from 0 (pure Mmd, in blue) to 1 (pure Mmm, in red). “Body condition” residuals along the hybrid index (for *Eimeria* spp. (b) and pinworms (c)) show no difference for infected mice (light green) and non-infected mice (grey). The optimized fit is represented by a solid line, the 95%CI of the fit as all parameters are allowed to vary in their 95%CI, is plotted as a grey ribbon. The 95%CI of the hybridization parameter alpha, as all parameters are fixed to their fitted value while alpha is allowed to vary in its 95%CI, is plotted as dashed lines.

## Discussion

We found lower intensities of the intracellular parasites *Eimeria* spp. and intestinal parasite pinworms in hybrid than in parental subspecies hosts in a previously unstudied transect of the European HMHZ. Lower intensity in hybrids is unlikely to be explained by epidemiological differences across the HMHZ, as we did not find the probability of infection to be similarly reduced in hybrid hosts.

House mouse hybrids are late generation in the European HMHZ (Macholán et al. 2007) and therefore should not be considered in categories, but rather on a continuous scale when analysing parasite infections or any other trait (Baird et al. 2012). We followed the statistical analysis of Baird et al. (2012) and explicitly modelled the effect of hybridization on parasite intensity by approximating the number of new combinations of genes brought together in a hybrid genotype by its expected heterozygosity (He). In other words we used He to derive non-linear predictions for hybridization effect based on the observed individual hybrid indices. This involved extending the existing approach to de-confound the prevalence and intensity aspects of parasite load. To increase reproducibility, we make our analysis available in an R package (Balard and Heitlinger 2019). The package allows statistical modelling with distributions additional to the original negative binomial distribution for (worm) count data (Baird et al. 2012). This allowed us to model the intensity of *Eimeria* infections as measured by a recently established quantitative PCR (Ahmed et al. 2019, Jarquín-Díaz et al. 2019, Al-khlifeh et al. 2019).

To our knowledge no studies have previously tested the effect of mouse hybridization on parasites other than helminths in a field setting of the HMHZ. To understand parasite processes in host hybrid zones, it is necessary to sample across different axes, one of them being pathogenicity of the parasite. *Eimeria* is very likely more pathogenic than pinworms (Al-khlifeh et al. 2019, Hakkarainen et al. 2007, Fuller and Blaustein 1996). The latter are common in laboratory facilities and often considered to provoke subclinical symptoms (Baker 1998) while mice experimentally infected by both *E. falciformis* and *E. ferrisi* have shown weight loss and diarrhoea (Al-khlifeh et al. 2019). Yet the pattern of reduced load in hybrid hosts is the same for the two parasites. These findings confirm that reduction in parasite intensity is either an effect intrinsic to the host individuals (e.g. enhanced immune reactions leading to increased resistance), or, if dependent on the parasite and/or parasite-host interplay, can be generalized over very different parasite species with different pathogenicity.

Adding more evidence to the original observation of reduced parasite loads for previously investigated parasites, we also found reduced pinworms loads in hybrids of our novel transect of the HMHZ. Despite some differences between the Brandenburg and Czech-Bavarian transects in pinworm infection such as distinct loads between males and females and lower prevalence (52.5%) and abundance (18.7) in the former compared to the latter (no significant difference between sexes; prevalence 70.9%, abundance 39.18; Baird et al. 2012), both the direction and strength of the hybridization effect were very similar in the two study areas. Since in various portions of the HMHZ there may be different ecological and epidemiological conditions this similarity reinforces our confidence that reduced parasite load in hybrids is intrinsic to the individual host or host-parasite interplay rather than a by-product of epidemiology.

A novel aspect of our work compared to previous studies of parasitism in the HMHZ is the separate study of parasite prevalence and intensity. This approach should not only reduce problems in statistical inference caused by false negative measurements (so called zero-inflation) but also allows us to address two different questions separately: (i) Is the *probability* of infection different for hybrids and pure subspecies? and (ii): Is there a difference in parasite *intensity* between infected hybrid and infected pure individuals?

An illustrative example of an ecological factor that could potentially lead to epidemiological differences is the density of hosts. Densities of mouse populations in the HMHZ centre may be lower than outside (either due to selection against hybrids or because the HMHZ as a tension zone tends to be trapped in “density troughs” sensu Hewitt 1975). Host density is expected to be positively correlated with pathogen transmission (Anderson and May 1979) and as a result prevalence may be higher in more dense populations (Morand and Guégan 2000). This is, however, not a general law as host density and *Eimeria* spp. prevalence are, for example, negatively correlated in bank voles (Winternitz et al. 2012). Independent of the direction of the effect, correlation between abundance and prevalence could be confounded with intrinsic effects of hybrid hosts.

Our analysis of prevalence (presence/absence in a logistic regression), did not however show any significant decrease of this probability of infection towards the centre of the zone, for neither *Eimeria* spp. nor pinworms. We argue here that, in conjunction with higher intensities, this distinguishes intrinsic hybrid effects from potential ecological and epidemiological confounders.

Animals tolerant of low-pathogenic parasites might not suffer fitness reduction during high parasitemia. This could be the case, for example, if the parasite is beneficial for the host’s interaction with other parasites (Heitlinger et al. 2017) or if immune responses against the parasite are costly relative to the harm it causes (Råberg et al. 2007). In addition, according to the “Old Friend” (or “Hygiene”) hypothesis, the constant presence of helminths in natural populations has led to the evolution of a background basal release of regulatory cytokines (Rook, 2009) which might in turn impact the outcome of more pathogenic infections. Even for relatively pathogenic parasites, such as *Eimeria,* differences in resistance could be uncoupled from health effects by differences in tolerance (Råberg et al. 2007). For these reasons parasite load in itself should not be blindly considered as a proxy for host health and certainly not for host fitness comparisons across hybrid zones (Baird and Goüy de Bellocq, 2019). We here used body condition as a proxy for the health component of host fitness. We, however, did not find evidence for differences in body condition between hybrids and pure mice upon infection. We conclude that we do not have evidence that lower parasitemia in hybrids increases their health.

It is worthwhile expanding on the above. Intensity of a particular parasite infection is not necessarily correlated with reduced health and a fitness decrease. For example, the fitness of sterile hybrids (always zero) is invariant to infection intensity. Moreover a hybrid host could be robust due to heterosis (though it may still be sterile). Even if we had found increased health of hybrids, this would not be interpretable as leading to a higher total hybrid fitness, as the parasite mediated health fitness component is only one (likely minor) component of overall fitness. It has been shown for example that male mice in the HMHZ centre have reduced fertility compared to parental individuals (Albrechtová et al. 2012, Turner et al. 2012). If reduced parasite intensity is host driven (and not a result of host-parasite interactions) one could conclude that some physiological systems (e.g. reproductive) may be more dependent on “co-adapted complexes”, while others – such as the immune system – benefit from diversity. This latter would be hybrid vigour in the narrow sense (Baird et. al. 2012), but would still not necessarily lead to any effects on host species barriers (Baird and Goüy de Bellocq, 2019).

We can in future ask whether host (immunity and resistance) parasite (infectivity and virulence) or their interactions are underlying reduced parasite intensity in hybrid house mice. *Eimeria* spp. are suitable pathogens to perform experiments and field studies in this endeavour.

A prime candidate locus for mediating a positive effect of hybridization on the immune system (hybrid vigour) is the major histocompatibility complex (MHC). In mice two genes of the MHC showed different levels of polymorphism as well as population structure with many alleles inferred to be shared between the subspecies by maintenance of ancestral polymorphism (Čížková et al. 2011). Additionally, the small demes of house mice can function as reservoirs of MHC alleles, contributing to the diversity of this system across demes and populations (Linnenbrink et al. 2018). The genetic structure of the MHC and especially polymorphism shared across subspecies should make these loci good candidates to investigate for mechanisms behind hybrid vigour, among a number of other loci including Toll-like receptors (Skevaki et al. 2015). Previous work on toll-like receptor 4 already suggests different evolutionary patterns between the house mouse subspecies (Fornuskova et al. 2014). For host parasite interactions major candidate loci are immunity related GTPases on the host side and rhoptry kinases in coccidia (Lilue et al. 2013).

Hybridization has played a significant role during and after the divergence of house mouse subspecies as well as during the formation of “classical inbred strains” (Yang et al. 2011). Improving our understanding of parasite process across the HMHZ provides valuable information on the house mouse as the model species with the most thoroughly understood immune system. A transfer of knowledge from this model might further understanding of the interplay between parasites and hybridizing species, our own as well as species relevant for conservation.

## Aknowledgements

Collection of data would not have been possible without the collaboration of farmers and house owners where mice were trapped. We also thank the students of bachelor of Biology of the Humboldt University of Berlin for their contribution in data collection.

## Additional files

### Supporting information

**Supplementary Table S1. Hybrid indexes, georeference and parasite load for each individual**

**Supplementary Table S2. Full parametrisation of the fitted models and comparison by G-test of the four nested hypotheses.** H0: same expected load for the subspecies and between sexes; H1: same expected load across sexes, but can differ across subspecies; H2: same expected load across subspecies, but can differ between the sexes; H3: expected load can differ both across subspecies and between sexes.

