## Supplementary table S2 for "Intensity of infection with intracellular *Eimeria* spp. and pinworms is reduced in hybrid mice compared to parental subspecies"

| Weibull distribution | Alpha (hybridization effect) (♀) | P-value (alpha vs no alpha) (♀) | Alpha (hybridization effect) ♂ | P-value (alpha vs no alpha) ♂ | L1 (load Mmd) (♀) | L1 (load Mmd) ♂ | L2 (load Mmm) (♀) | L2 (load Mmm) ♂ | S (shape) (♀) | S (shape) ♂ | G-test vs H0 | | | G-test vs H1 | | | G-test vs H2 | | |
|---|---|---|---|---|---|---|---|---|---|---|---|---|---|---|---|---|---|---|---|
| Eimeria intensit | | | | | | | | | | | dLL | dDF | p-value | dLL | dDF | p-value | dLL | dDF | p-value |
| **H0** | **0.74** | **0.02** | | | **-0.70** | | | | **2.33** | | | | | | | | | | |
| H1 | 0.85 | 0.01 | | | -1.01 | | 0.10 | | 2.38 | | 0.65 | 1 | 0.26 | | | | | | |
| H2 | 0.79 | 0.03 | 0.67 | 0.38 | -0.35 | -1.10 | | | 2.39 | 2.27 | 0.30 | 3 | 0.89 | | | | | | |
| H3 | 0.92 | 0.01 | 0.73 | 0.36 | -0.88 | -1.18 | 0.86 | -0.79 | 2.48 | 2.28 | | | | 0.49 | 4 | 0.91 | 0.83 | 2 | 0.43 |

| | Alpha (hybridization effect) (♀) | P-value (alpha vs no alpha) (♀) | Alpha (hybridization effect) ♂ | P-value (alpha vs no alpha) ♂ | L1 (load Mmd) (♀) | L1 (load Mmd) ♂ | L2 (load Mmm) (♀) | L2 (load Mmm) ♂ | A1 (aggregation Mmd) (♀) | A1 (aggregation Mmd) ♂ | A2 (aggregation Mmm) (♀) | A2 (aggregation Mmm) ♂ | Z (♀) (Deviation of aggregation from additive model) | Z ♂ (Deviation of aggregation from additive model) | G-test vs H0 | | | G-test vs H1 | | | G-test vs H2 | | |
|---|---|---|---|---|---|---|---|---|---|---|---|---|---|---|---|---|---|---|---|---|---|---|---|
| | | | | | | | | | | | | | | | dLL | dDF | p-value | dLL | dDF | p-value | dLL | dDF | p-value |
| H0 | 0.91 | 0.01 | | | 44.46 | | | | 1.78 | | | | -0.90 | | | | | | | | | | |
| H1 | 1.11 | < 0.001 | | | 32.12 | | 61.95 | | 1.75 | | 1.68 | | -0.77 | | 5.56 | 2 | <0.01 | | | | | | |
| H2 | 0.64 | 0.22 | 1.39 | < 0.01 | 49.76 | 39.60 | | | 1.72 | 1.88 | | | -0.73 | -1.79 | 7.72 | 4 | <0.01 | | | | | | |
| **H3** | **0.91** | **0.04** | **1.46** | **< 0.001** | **35.57** | **30.38** | **68.67** | **51.84** | **1.45** | **2.10** | **2.00** | **1.33** | **-1.04** | **-1.23** | | | | 9.01 | 6 | <0.01 | 6.85 | 4 | <0.01 |
